# Macrophages induce stromal differentiation and endothelialization in iPSC-derived kidney organoids

**DOI:** 10.1101/2025.11.13.688016

**Authors:** Ekaterina Pecksen, Marc Vives Enrich, Mi-Sun Jang, Sergey Tkachuk, Tamar Kapanadze, Cory P. Johnson, Florian P. Limbourg, Nico Lachmann, Kai M. Schmidt-Ott, Hermann Haller, Yulia Kiyan

## Abstract

Human induced pluripotent stem cell (iPSC) - derived kidney organoids are powerful models of kidney development and disease but lack mature vasculature. Here, we show that co-culture with human monocyte-derived macrophages (MDM) induces robust endothelial and stromal differentiation in iPSC-derived kidney organoids. MDM promoted the formation of lumenized capillary networks, an effect reproduced by iPSC-derived macrophages and the THP-1 monocyte line. In contrast, organoids without MDM displayed only transient endothelial differentiation that coincided with transient VEGFA upregulation and that regressed during maturation. MDM counteracted this transient phase by releasing soluble Neuropilin-1 (sNRP1), which sequestered VEGFA and stabilized late endothelial development, evidenced by sustained CD31 promoter activity. Transcriptomic profiling revealed that MDM redirected mesodermal fate suppressing lateral plate and cardiac mesoderm while promoting paraxial and intermediate mesoderm and the generation of FoxD1⁺ stromal progenitors via CXCL5 (ENA-78) secretion. These stromal cells supported late endothelial maturation and enhanced overall organoid differentiation. Our findings uncover a macrophage-driven mechanism coupling stromal and vascular development, providing a strategy to achieve functional vascularization in kidney organoids and improving their physiological relevance for research applications.

## Introduction

iPSC-derived organoids provide valuable models for disease, drug discovery, and regenerative medicine^1^. They replicate the structure and function of human organs, enabling investigation of more complex biological processes, compared to traditional 2D cell cultures. Kidney development relies on signaling crosstalk between the developing ureteric bud and metanephric mesenchyme. Key signaling molecules involved in this process have been identified, enabling the in vitro differentiation of stem cell–derived kidney organoids^2–4^. Further strategies implied separate differentiation of metanephric mesenchyme, ureteric progenitor cells, and stroma; these components were then combined to achieve higher structural maturation of the organoids/assembloids^5–7^. This strategy has allowed for the generation of more complex and physiologically relevant organoid structures, improving the fidelity and functionality of kidney models in vitro. However, many factors influencing kidney development remain unknown, which results in limited maturation of these organoids—particularly in terms of vascularization and functional capacity (filtration)^8^. The human kidney is highly vascularized, with blood vessels comprising more than half its structure. In the absence of vascularization, the biological and translational relevance of kidney organoids is questionable, rendering them effectively only a partial organ model. Furthermore, lack of vascularization is general problems in the organoid field.

Endothelial progenitors form during kidney-organoid differentiation in vitro by different protocols ^2,3,5,9^. However, capillaries show low density and complexity, do not associate with renal epithelium, and regress over time^10^. Since organoid differentiation in vitro follows an embryonic developmental program, the origin of renal endothelium in vivo must be addressed. Kidney vascularization is regarded to evolve by the combination of angiogenesis from pre-existing larger blood vessels and in situ vascular genesis from local endothelial progenitors^11,12^. However, the nature of these vascular progenitors remains unclear. Mugford and colleagues implicated Osr1-positive intermediate mesoderm progenitor cells as a source of endothelium^13^. Sims-Lucas showed that in mouse kidney FoxD1-positive stromal cells co-express endothelial markers^14^. They also demonstrated that isolated FoxD1-positive cells show endothelial and angiogenic properties in vitro such as tube formation and acetylated low-density lipoprotein uptake. Furthermore, conditional deletion of Flk1 in the FoxD1 lineage impaired peritubular capillary formation and renal morphology, suggesting a possible contributory role in specific endothelial compartments^15^. In vitro, Low et al. identified a subset of nephron progenitor cells as a source of endothelial cells during differentiation of iPSC-kidney organoids^16^.

Our earlier research showed that MDM could promote development of kidney organoids from iPSC^17^. Munro et al. showed that during murine kidney development, macrophages closely associate with forming vasculature and exhibit a pro-angiogenic phenotype^18^. Hence, we hypothesized that macrophages can promote vascularization of iPSC-kidney organoids and set out to test that notion.

## Results

### Myeloid cells promote endothelial differentiation in iPSC-kidney organoids

We reported recently that kidney-organoid differentiation was improved by co-culturing with human monocytes and iPSC-macrophages^17^. We further characterized the organoids developed in co-culture with myeloid cells and observed that not only were organoids better developed in the presence of macrophages^17^, but also differentiation of endothelial cells was strongly augmented. TaqMan RT-PCR performed after co-culturing iPSC with human monocytes showed greatly increased expression of endothelial markers on day 28 (Figure 1A) and advanced formation of the capillary network surrounding organoids (Figure 1B). 3D confocal microscopy showed that in co-culture endothelial cells were forming below the organoid and rose along the developing tubules towards the differentiating glomeruli (Figure 1C, Supplementary video 1 and 2). Higher magnification images disclosed extensive EC sprouting around the glomeruli (Figure 1D, Supplementary video 3). These results indicated better maturation of endothelial cells in co-culture. We also detected capillaries with lumen using 3D confocal microscopy (Figure 1E). Similar increases in endothelial differentiation were observed when iPSC were co-cultured with the monocytic cell line THP-1 (Figure 1F).

**Figure 1.**
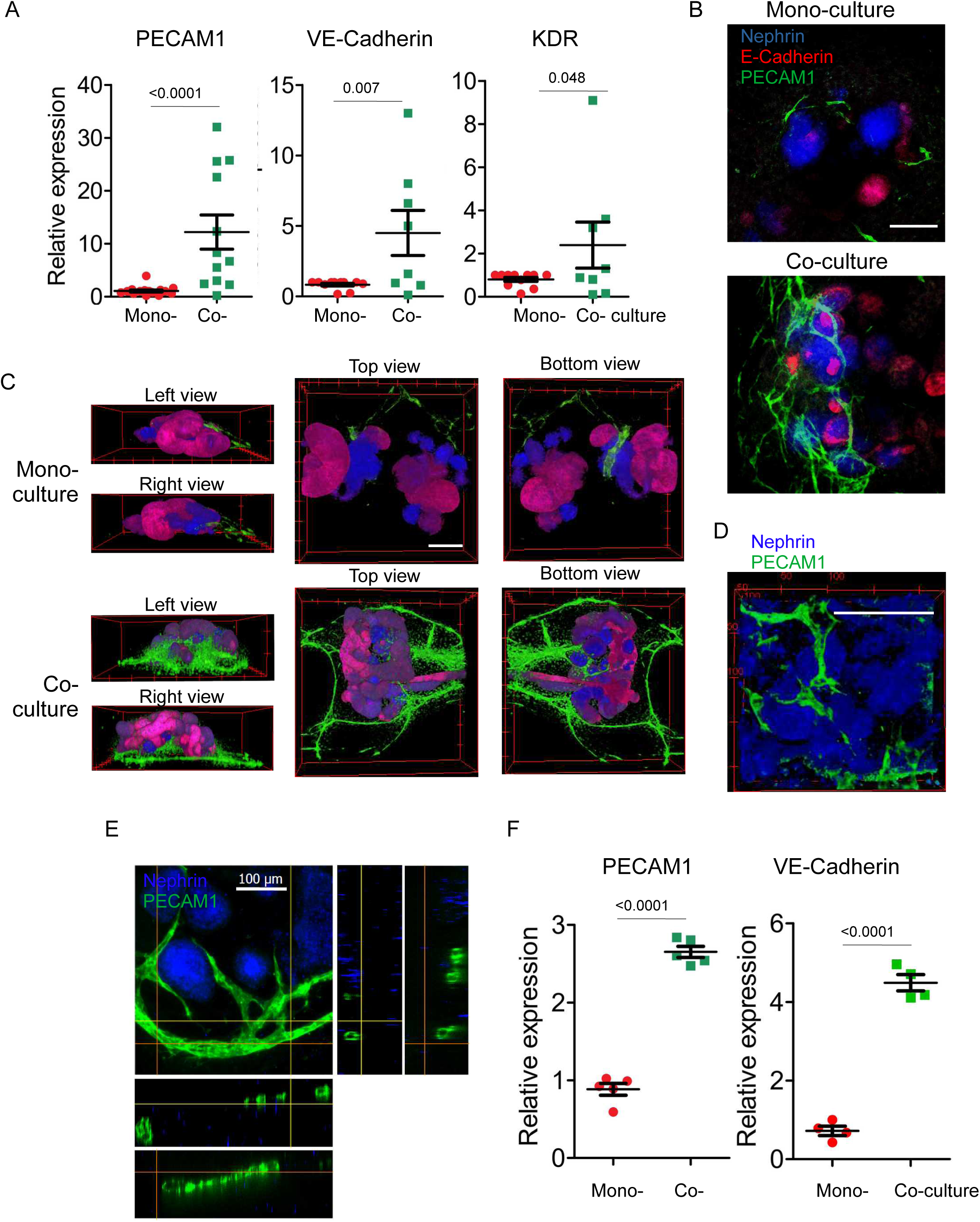
Myeloid cells promote endothelial differentiation in kidney organoids. A. Expression of endothelial markers was analyzed in kidney organoids differentiated from Epi-iPSC in mono-culture and in co-culture with MDM from day 0 until day 28 using TaqMan RT-PCR on day 28. B. iPSC-kidney organoids were fixed on day 28 and stained for Nephrin1 (blue), E-Cadherin (red) and PECAM1 (green). 3D confocal image stacks were acquired, and the sum projection of the stacks was generated using ImageJ. Scale bar is 100 µm. C. 3D confocal image stacks of organoids stained for Nephrin1 (blue), E-Cadherin (red) and PECAM1 (green) were processed using 3D-Viewer plugin of ImageJ. Scale bar is 100 µm. D. 3D confocal image stacks of organoids differentiated from Epi-iPSC co-cultured with THP-1 cells and stained for Nephrin1 (blue) and PECAM1 (green) were processed using 3D-Viewer plugin of ImageJ. Scale bar is 100 µm E. iPSC-kidney organoid was differentiated in co-culture with monocytes and stained for Nephrin (blue) and PECAM1 (green). Orthogonal views from 3D confocal image stacks are shown. Yellow and red lines indicate the corresponding positions of the orthogonal views. F. Expression of endothelial markers in iPSC kidney organoids differentiated in monoculture and in co-culture with THP-1 cell line from day 0 until day 28 was assessed by TaqMan RT-PCR on day 28.

### Monocytes differentiate into M2-type MDM

Monocytes are short-lived cells and in vitro they die rapidly unless macrophage differentiation is induced; we used FACS analysis to assess monocyte viability. We observed that the viability of MDMs declined gradually starting from day 8, both in mono-and co-culture with iPSC (Figure 2A) leading to >60% of cell death by day 14. Since such lengthy survival of monocytes without differentiation is unlikely, we characterized the phenotype of monocytes and iPSC-derived macrophages during the co-culture. Freshly isolated human classical monocytes represent a homogeneous CD14+/CD16-cell population (Supplementary Figure 1A). They rapidly changed their morphology, demonstrating an increase in granularity (Supplementary Figure 1A), differentiated into MDMs, and acquired macrophage-marker CD68, CD16, CD11c, HLA-DR and MerTK expression. This expression pattern is similar when the cells are treated with the same factors when co-cultured with iPSC even in the absence of the latter (Supplementary Figure 1C). The expression levels of CD206 and CD163 were significantly higher than CD86 suggesting that this environment promotes a shift in the MDM phenotype towards an M2 anti-inflammatory phenotype with a slightly more pronounced trend in co-culture with iPSC (Figure 2B-D). Since no M-CSF was added to the differentiation medium, we tested its expression by monocytes themselves. Monocytes expressed M-CSF and thus could promote differentiation to MDM in auto/paracrine fashion (Supplementary Figure 1D). Myeloid cell line THP-1 and iPSC-derived macrophages already expressed macrophage markers at the beginning of differentiation and did not demonstrate significant changes in the marker’s expression (Supplementary Figure 1E). The cell-survival kinetics were also similar to MDM.

**Figure 2.**
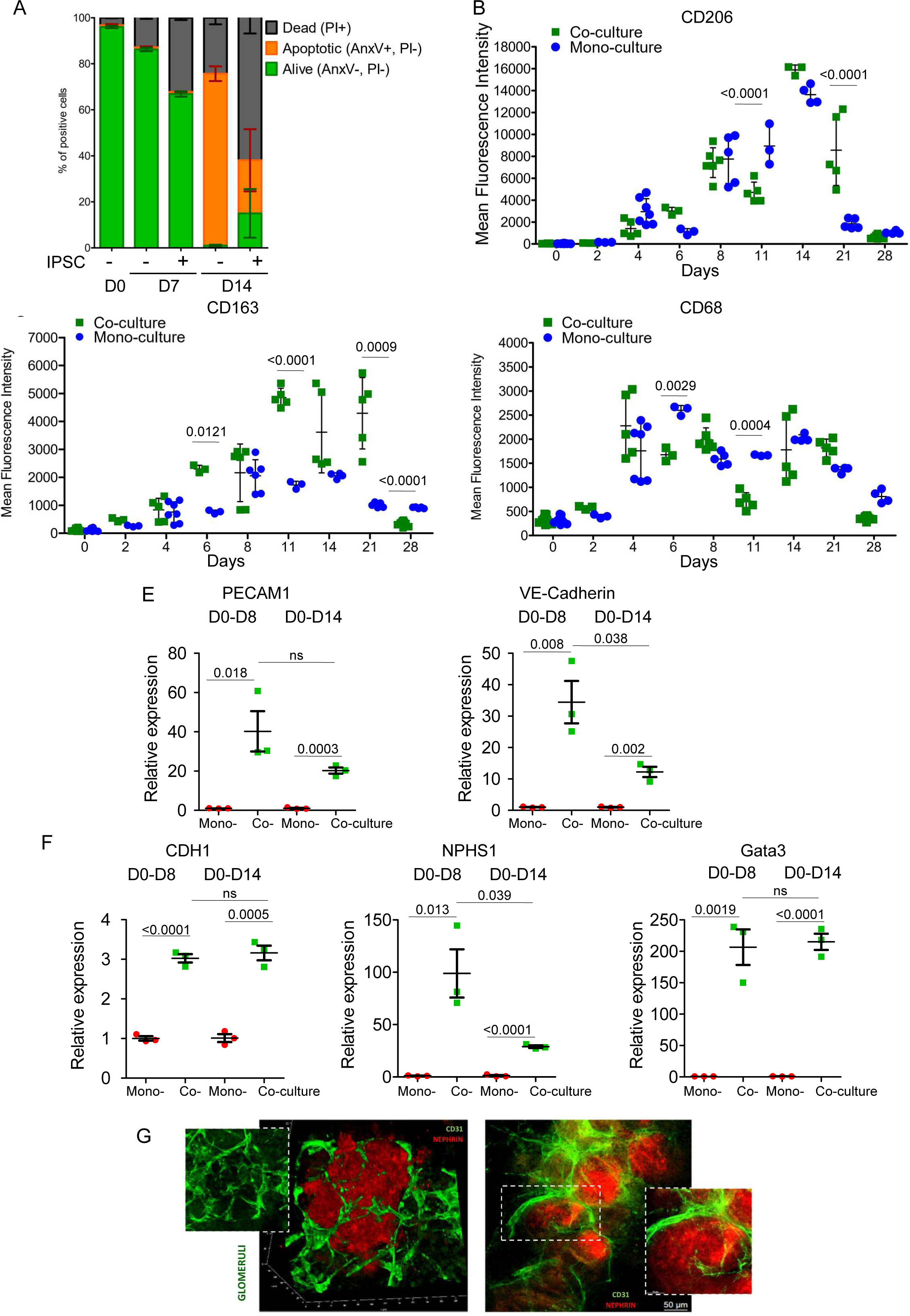
Monocytes co-cultured with iPSC kidney organoids differentiate to MDMs. A. Survival of monocytes in co-culture with iPSC was assessed by PI/AnxV staining and FACS analysis. B-D. Expression of CD206, CD163 and CD68 on MDMs was characterized by FACS analysis. MFI is shown. Expression of endothelial (E) and renal (F) markers on kidney organoids differentiated from Epi-iPSC co-cultured with MDMs from day 0 (D0) until day 8 (D8) and from day 0 (D0) until day 14 (D14) was assessed by TaqMan RT-PCR. G. cl11/Selene iPSC were differentiated to kidney organoids in co-culture with iPSC-derived macrophages day 0 until day 8 (right image) and from day 0 until day 14 (left image).

Based on the viability kinetics of MDM, we assumed that their window of action should be within the first two weeks of the iPSC differentiation and performed co-culture with MDM from day 0 until day 8 and from day 0 until day 14. The strongest impact on the differentiation of organoids and endothelial cells was observed by co-culturing iPSC with MDMs until day 8 (Figure 2E, F). Co-culturing until day 14 was less efficient. As analysis of MDMs survival showed a significant number of apoptotic cells on day 7 (Figure 2A), these data suggested that their effects began to decline afterward and the presence of apoptotic macrophages can partially reverse their positive influence. These observations were confirmed by co-culturing cl11/Selene iPSC with iPSC-derived macrophages from day 0 until day 8 and from day 0 until day 14 (Figure 2G) using 3D confocal microscopy.

### Endothelial cells form capillary network

We used FACS analysis to quantitatively assess endothelial cell population in mono-and co-culture. Population of CD31-positive cells was strongly increased in co-culture compared to monoculture (Figure 3A). Endothelial cells were also positive for VE-Cadherin (CD144) and CXCR4 (Figure 3B). Further, we applied 3D confocal microscopy to visualize and characterize EC vascular network surrounding individual organoids. Tile scanning of the total area of over 2 mm^2^ revealed a capillary network interconnecting adjacent organoids (Figure 3C, Supplementary Figure 2). To characterize the capillary network, we quantified total vessel area, total number of end points, total number of junctions and vessel lacunarity using AngioTool software^19^. All these analyses provided a consistent outcome of an increased number of endothelial cells and better organized vascular network in the organoids co-cultured with MDM (Figure 3D-H).

**Figure 3.**
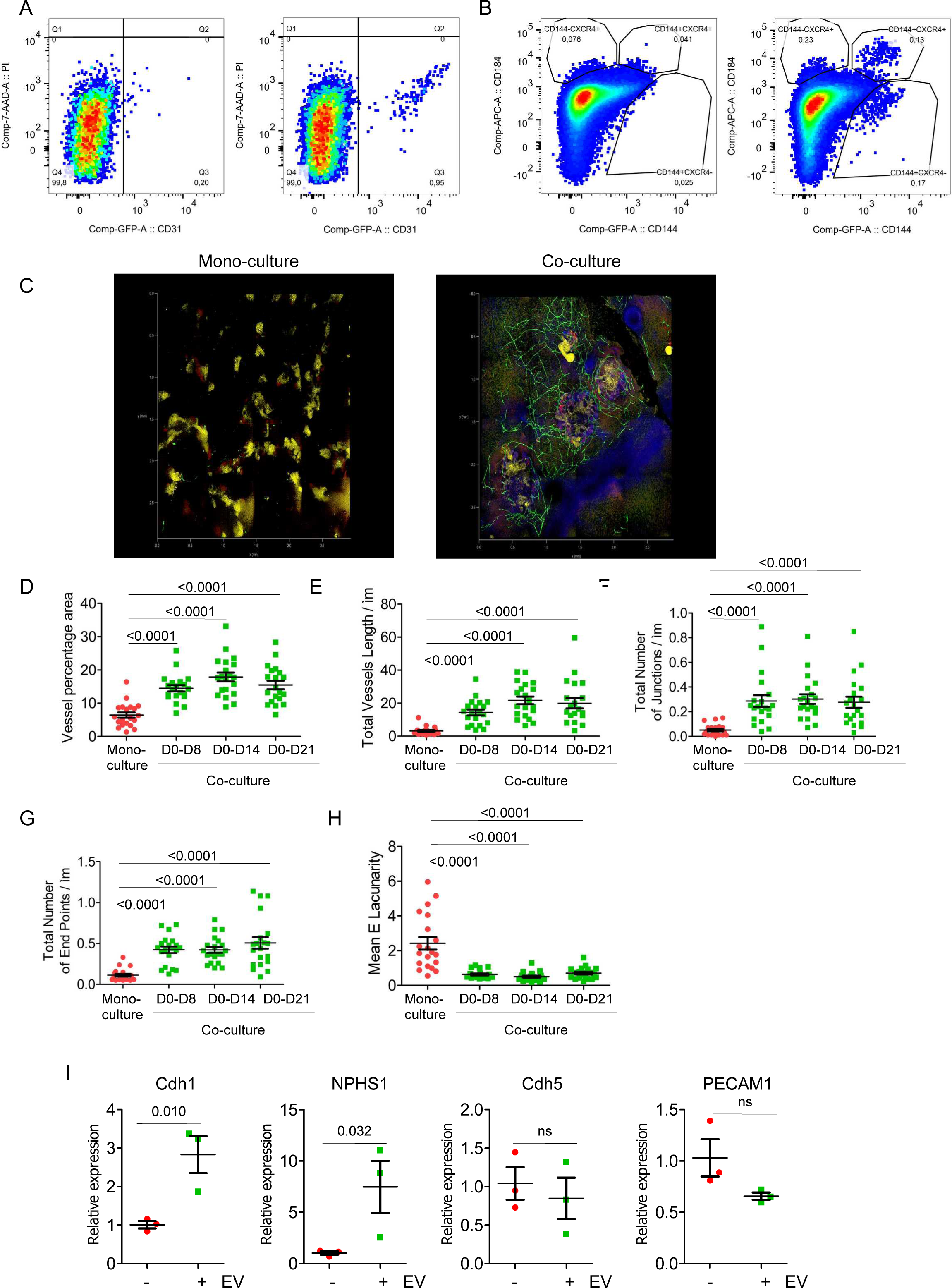
Macrophages promote capillary network formation in iPSC-kidney organoids. A. Organoids differentiated from Epi-iPSC in monoculture and in co-culture with MDM were enzymatically digested and endothelial cells were analyzed by FACS. B. Tile scan of 2 mm² area of cl11/Selene iPSC differentiated to kidney organoids in monoculture (left) and in co-culture with iPSC-derived macrophages (right) from day 0 until day 8. Organoids were stained and fixed on day 28. C-G capillary network was characterized using AngioTool. Vessel percentage area (C), Total vessel length (D), Total number of junction (E), Total number of end points (F) and Mean lacunarity (G) of capillary networks formed in monoculture or in co-culture with iPSC-derived macrophages from day 0 (D0) until day 8 (D8), day 14 (D14) or day 21 (D21) were analyzed. H. Epi-iPSC were differentiated to kidney organoids with or without addition of MDM-derived EV. EV were added from day 0 until day 14. Expression of markers was analyzed by TaqMan RT-PCR on day 28.

We have previously shown that MDM promote iPSC survival during differentiation to kidney organoids via the release of extracellular vesicles (EV) ^17^. To test whether EV also promote endothelial differentiation, we treated monocultured MDM with the same medium as was used for iPSC differentiation, isolated EV by means of ultracentrifugation, and applied EV to monocultured iPSC from day 0 until day 14 during each medium change. On day 28 we performed RT-PCR. As shown in Figure 3J, EV promoted expression of renal markers Cdh1 and Nephrin, though Nephrin expression was lower than in co-culture with MDM. However, EV had no effect on endothelial differentiation. These data suggested that endothelial differentiation was rather induced by a soluble factor released by macrophages and not by EV.

### MDM prevent transient endothelial differentiation

MDM co-cultured with iPSC during the first 8 days of kidney organoid differentiation strongly promoted the development of organoids and induced differentiation of endothelium by day 28. To follow the kinetics of endothelial differentiation, we developed a CD31 promoter reporter cell line expressing secreted Gaussia luciferase. EF1alpha promoter was used as a negative control. Conditioned medium was collected during the differentiation of iPSC in mono- and co-culture and luciferase activity was measured. In monoculture, we observed a transient increase in CD31 promoter activity around day 14 followed by a decrease to the basal level by the end of the differentiation protocol. This result corresponds to the data reported by Cleaver et al.^10^ who also demonstrated the transient nature of endothelial differentiation during kidney organoid development. On the contrary, in co-culture with MDM, this transient increase was not observed, and activity of the CD31 promoter increased gradually during the organoid development (Figure 4A). Interestingly, CD31 promoter activity on day 14 was higher in monoculture than in co-culture. A transient increase in endothelial differentiation in monoculture was confirmed by immunocytochemistry (Figure 4B) and RT-PCR (Figure 4C).

**Figure 4.**
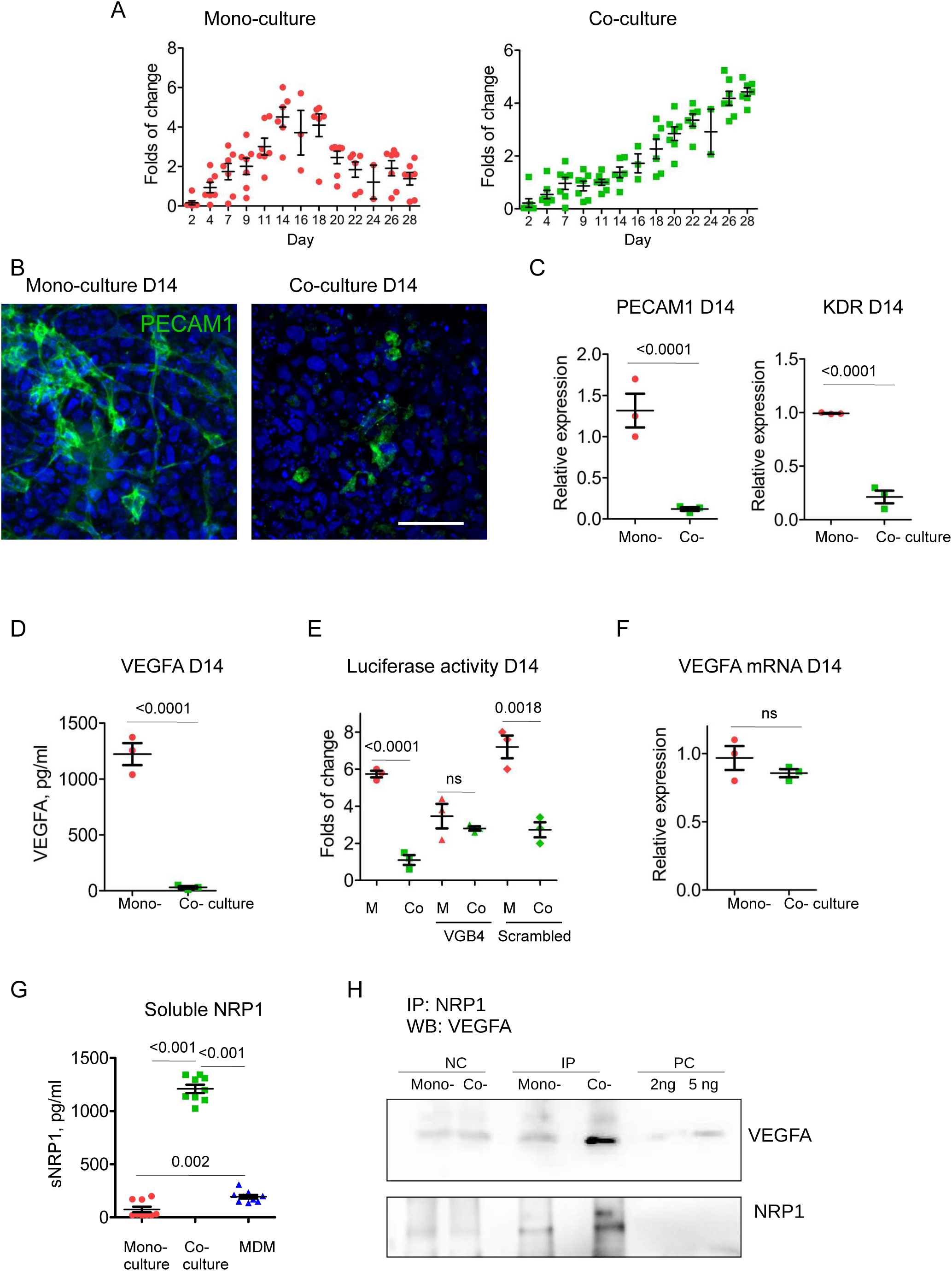
MDM suppress transient endothelial differentiation in iPSC-kidney organoids by antagonizing VEGFA. A. iPSC reporter cell line expressing Gaussia luciferase under PECAM1 promoter was differentiated to kidney organoids in mono-culture (left) and in co-culture (right) with MDM. Luciferase activity was measured in conditioned medium. B, C. Immunocytochemistry (B) and TaqMan RT-PCR (C) was performed in Epi-iPSC differentiating towards kidney organoids in mono- and co-culture with MDM on day 14. PECAM1 (green), DAPI (blue). Scale bar is 50 µm. 14. D. Concentration of VEGFA in conditioned medium was assessed on day 14 by ELISA. E. PECAM1-gaussia luciferase reporter cell line was differentiated to organoids in mono-culture and in co-culture with MDM. VGB4 or control scrambled peptides were added from day 0. Luciferase activity was assessed on day 14. F. VEGFA mRNA expression was measured by TaqMan on day 14. G. Concentration of sNRP1 was assessed in conditioned medium of mono-cultured Epi-iPSC, co-cultured with MDM and in medium of MDM treated with the same factors without iPSC on day 14. H. Association of sNRP with VEGFA in conditioned medium of mono- and co-cultured Epi-iPSC was evaluated by immunoprecipitation followed by western blotting.

Since VEGFA is the strongest factor of endothelial differentiation, we measured concentration of VEGFA on day 14 by ELISA. Concentration of free VEGFA was higher in monoculture in compared to co-culture (Figure 4D). Assuming that VEGFA is the driving factor of transient endothelial differentiation in monoculture, we assessed CD31 promoter activity in the presence of VGB4 peptide blocking VEGF/VEGFR interaction. Scrambled peptide was used as a control. Confirming our assumption, CD31 promoter activity was decreased by VGB4 peptide in monoculture, whereas this effect was not shown on day 14 in co-culture with MDM (Figure 4E). Since VEGFA mRNA expression by iPSC was not changed by co-culturing with MDM (Figure 4F), we hypothesized that MDM release a soluble factor that can antagonize VEGF/VEGFR signaling at this stage. Neuropilins 1 and 2 (NRP1 and NRP2) can bind VEGFA and act as co-receptors, promoting VEGFA signaling towards angiogenesis and beyond that^20^. However, when NRP1 is shed from the cell surface by protease, it can still bind VEGFA thus preventing its interaction with VEGF receptors on the cell surface^21^ and acting as a decoy receptor. Hence, we analyzed the concentration of soluble neuropilins in conditioned medium on day 8 by ELISA. NRP2 was not detected in soluble form, whereas concentration of NRP1 was very high in co-culture but not in monoculture of iPSC or MDM cultured alone (Figure 4G). To confirm the interaction of NRP1 and VEGF, we performed a co-immunoprecipitation assay and detected VEGFA co-immunoprecipitated with NRP1 from conditioned medium of iPSC co-cultured with MDMs (Figure 4H). These data suggest that VEGFA released by iPSC promoted transient endothelial differentiation by the auto/paracrine signaling. MDM released soluble NRP-1 and antagonized VEGF signaling and the development of transient endothelium.

### MDM orchestrate progenitor differentiation

The above data indicated that MDM affect kinetics of endothelial differentiation and argue for the non-identical nature of endothelial progenitors in mono- and co-culture. To gain an insight on how MDM affect transcriptional profile of iPSC, we performed bulk RNA-sequencing. We decided to perform RNA-sequencing of iPSC on day 8 because co-culturing of iPSC with MDM from day 0 until day 8 was the most efficient to induce endothelial differentiation detected on day 28. Though the strongest CD31 promoter activity in monoculture was observed on day 14, we reasoned that on day 8 the progenitor population of early transient endothelium in monoculture should already be present. Analysis of RNA-sequencing data revealed striking differences in the gene expression pattern of iPSC in mono- and co-culture (Supplementary Figure 3A-C). We identified 1505 upregulated and 1671 downregulated genes with p-adj<0.05 and log2(FC) values greater than 1 or below -1 (Supplementary Figure 3, Supplementary Table 2). Volcano plot showed among the most upregulated genes were MEOX2 involved in presomitic mesoderm differentiation, SIX1 important for the differentiation of intermediate mesoderm, and FOXD1, a marker of renal cortical stroma (Supplementary Figure 3B). We then analyzed marker expression of the three main mesoderm lineages (Figure 5A). MDM decreased differentiation of lateral/cardiac mesoderm, whereas specification of paraxial mesoderm, mesodermal derivatives, and intermediate mesoderm, were strongly promoted. This observation was confirmed by the enrichment analysis of downregulated genes, as main Gene Ontologies identified were related to Heart development (GO:0007507) (Figure 5B). In addition, our data on the inhibition of VEGF-dependent signaling by MDM, was confirmed. Expression of VEGFR2 (KDR) and VEGFR1 (FLT1) were downregulated in co-culture and Blood vessel development related Gene Ontologies were identified by enrichment analysis of downregulated genes (Supplementary Figure 3D). Pathways related to heart development and muscle differentiation are considered off-target processes during kidney organoid differentiation in vitro. In co-culture with MDM we observed very strong downregulation of genes related to those pathways. Expression of VEGFA and VEGFB remained unchanged by MDM. The full lists of GOs and genes are given in the Supplementary Table 3. Loh et al. stepwise mapped the differentiation of iPSC to different mesoderm cell types and showed that specification of lateral mesoderm is inhibited by WNT signaling^22^. Interestingly, MDM induced expression of several WNT ligands and receptors in iPSC and KEGG WNT signaling was identified by enrichment analysis of DEG (Figure 5C, D).

**Figure 5.**
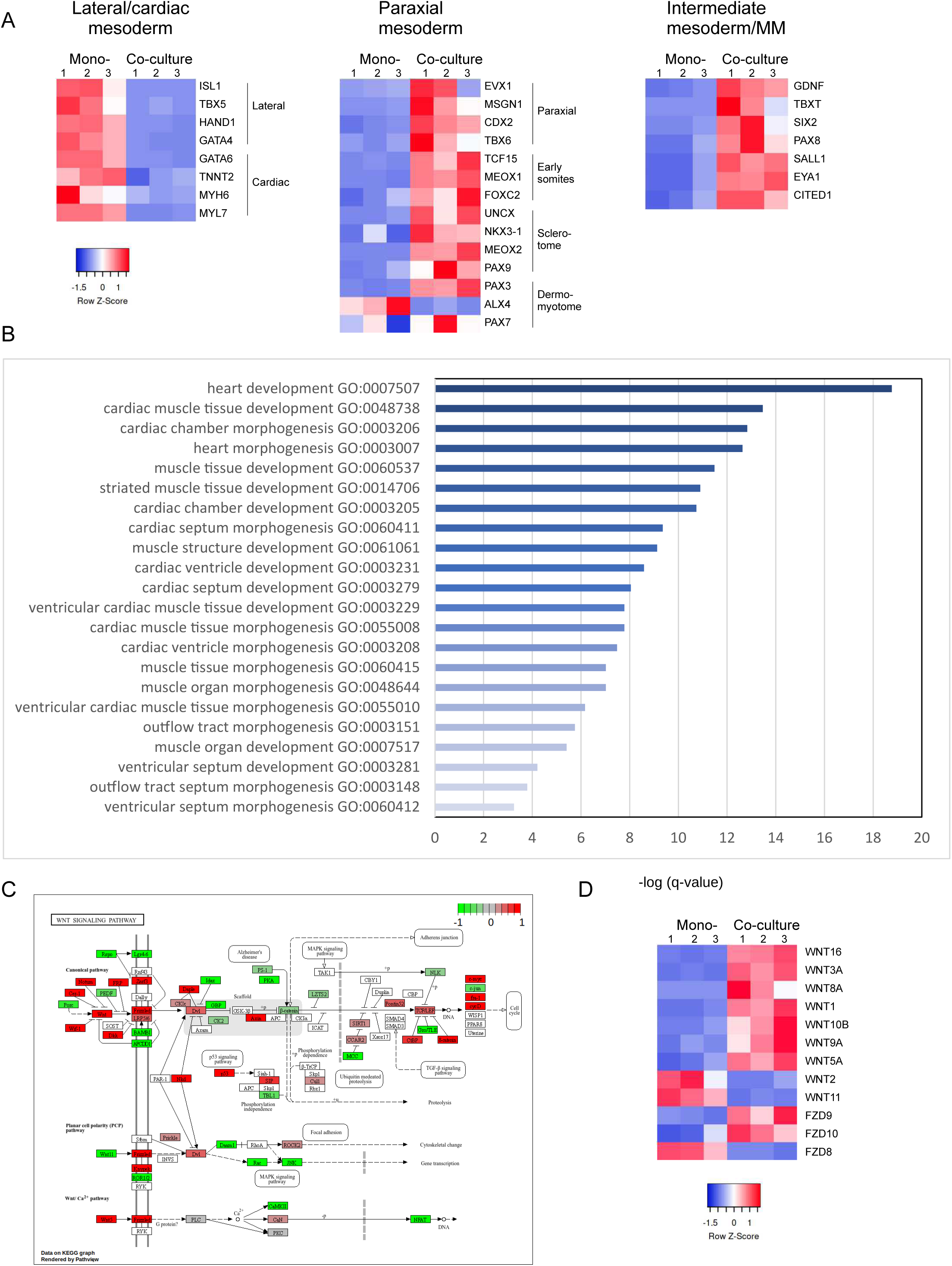
MDM orchestrate mesoderm patterning in differentiating iPSC. A. Heatmaps of Lateral plate/cardiac, paraxial and intermediate mesoderm/metanephric mesenchyme markers expression in mono- and co-cultured Epi-iPSC based on RNA-sequencing data. B. Most significantly enriched GO terms identified by enrichment analysis of DEG downregulated in co-cultured Epi-iPSC in comparison to mono-cultured. C. Visualization of WNT Signaling pathway identified by KEGG enrichment analysis with gene expression values mapped to gradient color scale. D. Heatmap of WNT ligands expression in mono-and co-cultured Epi-iPSC based on RNA-sequencing data.

Enrichment analysis of upregulated genes showed their relation to Tube morphogenesis (GO:0035239); Tissue morphogenesis (GO:0048729); and Pattern specification process (GO:0007389). Child terms of these three GOs and their -log(q-value) are displayed (Figure 6). Pathways upregulated in co-culture were strongly related to vascular and kidney development. However, the pathways did not contain genes related to VEGF/VEGFR signaling, except for upregulated VEGFD. Somitogenesis was also identified in the enrichment analysis, confirming the formation of paraxial mesoderm. Genes associated with identified GO are listed in the Supplementary Table 4.

**Figure 6.**
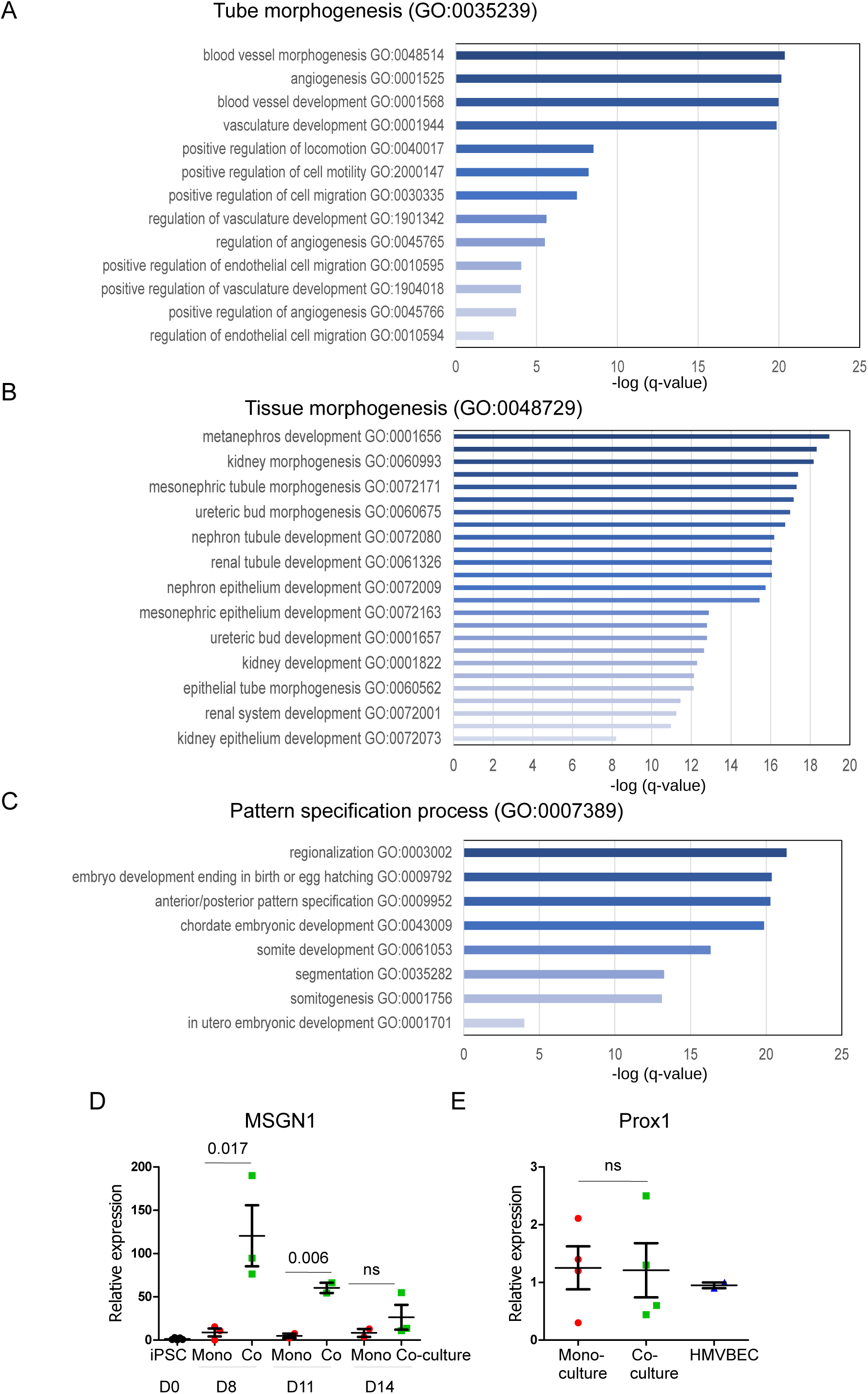
MDM promote vascular and renal development in differentiating iPSC. Child terms of the most significantly enriched GO terms identified by enrichment analysis of DEG upregulated in co-cultured Epi-iPSC in comparison to mono-cultured. A. Tube morphogenesis (GO:0035239). B. Tissue morphogenesis (GO:0048729). C. Pattern specification process (GO:0007389). D. Expression of MSGN1 was assessed by TaqMan in mono-cultured Epi-iPSC and co-cultured with MDM. E. Expression of PROX1 in CD31+ cells isolated from organoids differentiated in monoculture and in co-culture with MDM on day 28.

RNA-sequencing data suggested that MDM switch the differentiation mesoderm subtypes. Indeed, markers of lateral plate mesoderm and cardiac mesoderm were strongly downregulated by MDM (Figure 5A). Takasato et al.^23^ reported spontaneous generation of a lateral plate mesoderm from embryonic stem cell-derived posterior primitive streak. PDGFRA and Prrx1-expressing cells have also been identified as off-target cells during kidney organoid differentiation detected by single-cell RNA-sequencing^24^. Therefore, we were not surprised to find lateral plate mesoderm-fate committed cells as an off-target population. Furthermore, we demonstrated that MDM prevented this pathway of differentiation. On the contrary, markers of the paraxial mesoderm were strongly upregulated in co-culture (Figure 5A). Expression of posterior intermediate mesoderm and metanephric mesenchyme markers corresponding to day 8 of the protocol was also upregulated.

We also verified increased expression of MSGN1 by RT-PCR (Figure 6D). Confirming RNA-sequencing data, expression was strongly upregulated by MDM on day 8 and consequently decreased by day 14. It is known that during the development, paraxial mesoderm gives rise to the lymphatic vasculature^25^. To rule out whether endothelial cells in organoids co-cultured with MDM originate from induced paraxial mesoderm and have a lymphatic nature, we isolated CD31+ cells from mono- and co-cultured organoids and assessed expression of transcription factor responsible for the development of lymphatic vasculature Prox1. We did not observe increased expression of Prox1 in co-culture (Figure 6E). Furthermore, expression level was also similar to human primary blood capillary endothelial cells suggesting that endothelial cells in organoids co-cultured with MDM are not lymphatic in origin.

Earlier, we showed that MDM rescue iPSC from CHIR-induced apoptosis. Here, KEGG analysis of RNA-sequencing data revealed upregulation of cell cycle-related genes (Figure 7A) suggesting that MDM effects extended beyond decreasing cell death to promoting proliferation. Genes involved in Ribosome Biogenesis (Supplementary Figure 4A) were also upregulated indicating increase in proliferation and differentiation during gastrulation^26^. Accordingly, Nucleocytoplasmic Transport and splicing (Supplementary Figure 4B, C) were also increased. Together, these data showed that MDM suppress lateral mesoderm differentiation via increased WNT signaling and thus prevent development of transient endothelium in organoids. Furthermore, MDM direct iPSC to paraxial and intermediate mesoderm development.

**Figure 7.**
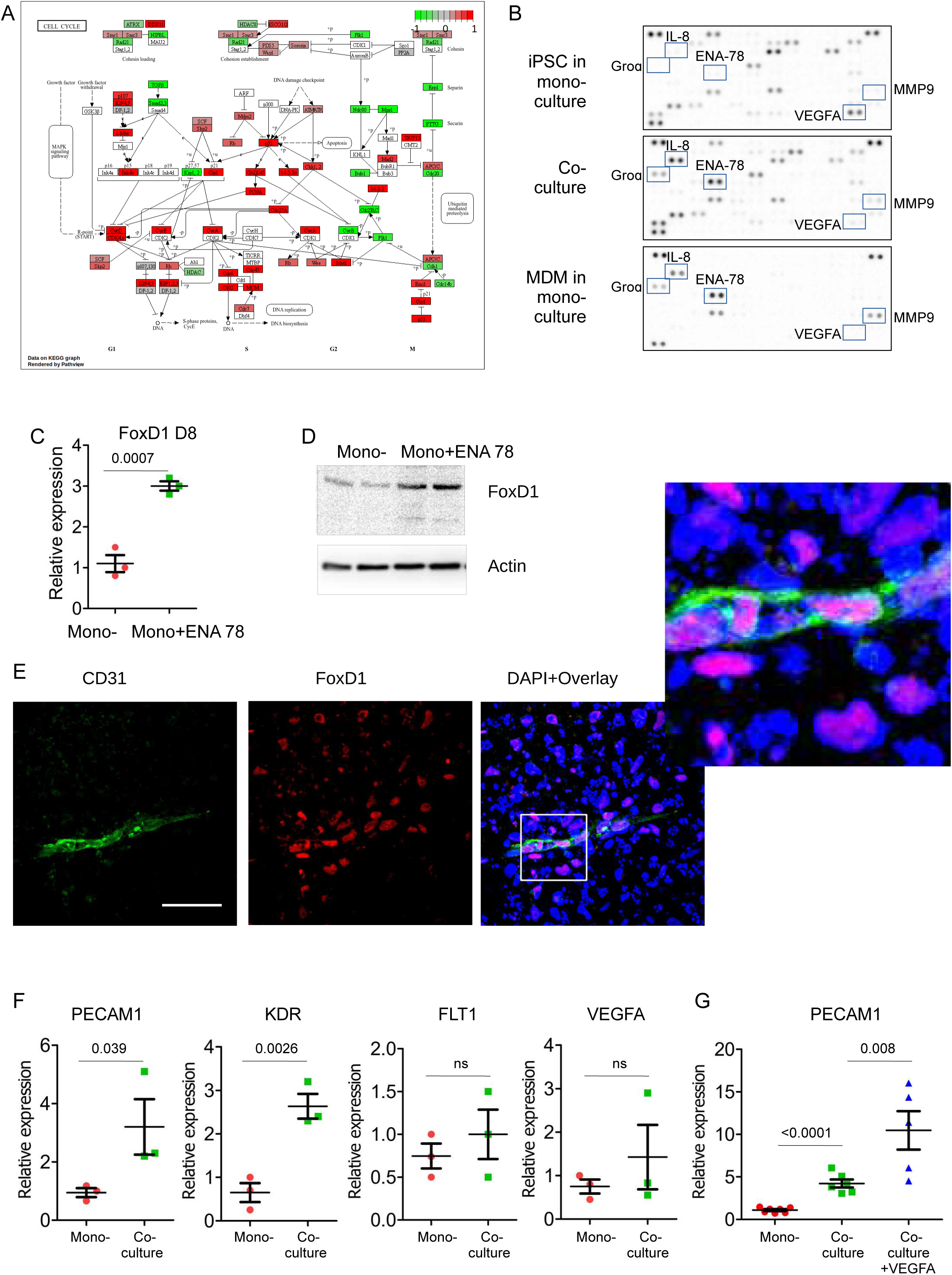
MDM promote FoxD1 stromal progenitor differentiation. A. Visualization of Cell cycle pathway identified by KEGG enrichment analysis with gene expression values mapped to gradient color scale. B. Proteome profiler human XL cytokine array was performed using conditioned medium from Epi-iPSC differentiating in mono- and co-culture with MDM. Most significantly altered cytokines are marked. C-D. FoxD1 expression in mono-cultured Epi-iPSC without and with addition of 15 ng/ml ENA-78 was evaluated on day 8 by TaqMan RT-PCR (C) and western blotting (D). E. FoxD1+ endothelial cells were detected on day 14 in Epi-iPSC co-cultured with MDM. Scale bar is 50 µm. F. Endothelial markers expression in CD31+ cells isolated from Epi-iPSC mono-cultured and co-cultured with MDM on day 28. G. Organoids co-cultured with MDM from day 0 until day 8 were treated with VEGFA from day 24 until day 28. PECAM1 expression was assessed by TaqMan RT-PCR.

### MDM induce FoxD1 expression via secretion of ENA-78

Since co-culturing of iPSC with MDM until day 8 was the most effective method to induce endothelial differentiation, we performed proteome profiling using human cytokine array membranes on day 8. We compared conditioned medium from mono-cultured iPSC, mono-cultured MDM and co-cultured iPSC with MDM (Figure 7B). The full list of cytokines is given in the Supplementary Table 1. Among the most highly expressed cytokines, expression of ENA-78, GROɑ, and IL-8 was strongly increased in co-culture. These cytokines were produced by MDM independently of the iPSC presence. However, extending differentiation protocol with IL-8 and GROα had no significant effect on organoid and endothelium differentiation. ENA-78 (CXCL5), on the contrary, was responsible for the upregulation of FoxD1 expression. FoxD1, a marker of renal cortical stroma, is important for kidney development. FoxD1-knockout mice demonstrate abnormal nephrogenesis and kidney hypodysplasia^27,28^. Furthermore, Sims-Lucas reported that FoxD1-positive stroma can give rise to endothelial cells and form a subset of peritubular capillaries^29^. Hence, strong increase of FoxD1 expression in co-culture with MDMs reflected the formation of stromal progenitors that can give rise to the development of late endothelium in co-culture with MDM. The role of CXCL5 in promoting cancer angiogenesis via FoxD1 induction has also been shown^30^. Interestingly, ENA-78 was among the most upregulated cytokines detected in co-culture with MDMs by Proteome Profiler Array (see Fig. 7B). To confirm this hypothesis, we stimulated iPSC differentiating to kidney organoids in monoculture with ENA-78 from day 5 till day 7 and observed increased expression of FoxD1 by TaqMan RT-PCR and western blotting (Fig. 7C, D). Furthermore, in co-culture with MDMs, a population of endothelial cells demonstrating FoxD1-positivity have been detected by immunocytochemistry (Figure 7E), suggesting that macrophages promote late endothelial differentiation in co-culture by antagonizing VEGFA/VEGFR signaling around day 14 and inducing FoxD1 stromal progenitors that give rise to later endothelial differentiation.

We were surprised that the late persistent vasculature in MDM-co-cultured organoids originated from antagonizing VEGF/VEGFR signaling on day 14. To better characterize endothelial cells, we isolated CD31+ cells from organoids on day 28 by means of magnetic beads and performed RT-PCR analysis for endothelial markers. Endothelial cells isolated from co-cultured organoids demonstrated higher expression of CD31 (PECAM1) and VEGFR2 (KDR), whereas expression of VEGFR1 and VEGFA was comparable in cell isolated from mono- and co-culture (Figure 7F). These data suggest that independently of their origin, late vasculature can be regulated by endogenous VEGF/VEGFR signaling. To confirm this, we induced organoid differentiation in co-culture with MDM from day 0 till day 8, then we withdrew MDM and added VEGF from day 24 until day 28. On day 28 we performed RT-PCR. As shown in Figure 7G, addition of VEGF further promoted development of endothelium in kidney organoids. Thus, vasculature induced by MDM demonstrates VEGFR expression and its development can be further promoted by VEGF during the late stage of organoid development.

## Discussion

Aiming to improve the development of iPSC-kidney organoids, we have tested the hypothesis that macrophages can influence their vascularization. We used three different iPSC cell lines and co-cultured them with several types of myeloid cells – MDM, iPSC-derived macrophages and THP-1 cell line - and demonstrated that myeloid cells change the landscape of iPSC differentiation resulting in strongly promoted endothelial differentiation. Interestingly, we were able to show that at least two mechanisms of vascularization exist in organoids: an endogenous, VEGF-dependent formation of endothelial cells, which is self-limited and short lived and the macrophage-dependent vascularization, which is stronger and persists. In the presence of macrophages, endothelial cells form a sophisticated vascular network that penetrates and interconnects organoids. 3D confocal microscopy showed that capillary form hollow lumen showing a high degree of cell specification. Inside the organoid, endothelial cells demonstrate intensive sprouting in glomerular differentiation. Quantitative characterization by means of AngioTool showed high complexity of the macrophage-dependent vascular capillary network (Figure 3).

Vasculature comprises an essential part of the kidney. However, stem cell-derived organoids fail to develop vascular system. Though endothelial cells were detected during kidney organoids differentiation by different protocols, their number was very low, and capillary network was not well-developed^10^. When transplanted into the mouse kidney or intracoelomic in chicken ^31^, organoids showed improved vascularization, however, most of endothelium was built up by the host cells. Several angiogenesis-promoting approaches have been used to improve endothelial development in kidney organoids. Overexpression of Hypoxia Induced Factor 1α (HIF1α)^32^ or placing the developing organoids in physiological hypoxia conditions^33^ promoted endothelial differentiation. Inducible overexpression of ETS translocation variant 2 (ETV2)^34^ also promoted endothelial development and improved multilineage maturation of the organoids^35^. Placing iPSC-derived kidney organoid under flow conditions has also improved vascular maturation^36^. However, those studies have not addressed what progenitors gave rise to the endothelium and it is unclear whether these approaches are relevant to the developmental program. Transplantation of recently reported kidney assembloids, that showed far more advanced structure than organoids have shown so far, demonstrated that their vascularization in vivo is to a significant degree formed by progenitors originating from the assembloids themselves^7^.

Not surprisingly, hypoxia and HIF1α -promoted vascularization relied on VEGF signaling. We have also showed that transient endothelial differentiation in mono-cultured iPSC was VEGF-dependent since it was abrogated by VGB4 peptide blocking VEGF/VEGFR interaction (Figure 7). Interestingly, MDM antagonized VEGFA signaling on day 14 via its binding with soluble form of NRP-1 (Figure 7). We confirmed this interaction by means of immunoprecipitation that was detected in co-culture with MDM but not in mono-cultured iPSC. By antagonizing VEGF action, expression of VEGFR2 (KDR) and VEGFR1 (FLT1) was downregulated. This resulted in prevention of transient increase in the differentiation of endothelial cells in the middle of the protocol in co-culture with macrophages. MDM-induced “late” endothelium derived from FoxD1-positive progenitors, as we detected a population of CD31+/FoxD1+ double-positive cells on day 14 of differentiation (Figure 7). Independently on the origin, vasculature developing under influence of MDM acquires VEGFR expression and becomes VEGF sensitive by day 28. This was confirmed by profiling CD31+ cells isolated on day 28 and adding VEGFA during the last four days of organoid development (Figure 7F, G).

Macrophages play a multifaceted role in angiogenesis, both direct and indirect, in vivo and in vitro. The angiogenesis-promoting role of tumor-associated macrophages is the most studied^37^. In vitro, macrophages also promoted vascularization of tissue-engineered scaffolds. Interestingly, pro-inflammatory M1-type macrophages were shown to secrete highest amount of VEGF, whereas M2a and M2c released PDGFBB and MMP9, correspondingly^38^. In addition to growth factors, pro-angiogenic role of macrophage-secreted EV has been reported^39^. We have previously showed that MDM-derived EV promote iPSC survival and organoid development^17^. However, we did not observe an increased endothelial differentiation when iPSC were treated with EV (Figure 3) suggesting the involvement of other MDM-derived factors in the differentiation and vascularization process. Having performed bulk RNA-sequencing on day 8, we observed that MDM have a stronger effect than just enhancing endothelial cell differentiation. We observed that genes of cardiac and lateral plate mesoderm were downregulated, whereas paraxial mesoderm and somitogenesis as well as intermediate mesoderm genes were strongly induced by MDM. This indicates that factors released by MDM affect the fate of iPSC during the specification of mesoderm.

How mesoderm specification is regulated and the role macrophages can play in this process is largely unknown^40^. Crosstalk between macrophages and surrounding embryonic tissues is also not well characterized and released factors remain largely unknown. Our model can serve to identify the role of macrophage-released factors during mesoderm specification. MDM induce WNT ligands expression in iPSC and thus can suppress differentiation of lateral mesoderm^22^. The fact that MDM induced specification of paraxial mesoderm and somitogenesis was unexpected. Other investigators reported earlier that paraxial mesoderm gives rise to lymphatic vascular system^25^. Therefore, we initially assumed a lymphatic nature of MDM-induced endothelium in our system. However, CD31-positive cells isolated from organoids co-cultured with MDM did not show increased expression of the main transcription factor responsible for lymphatic vasculature development, Prox1^41^ in comparison to monoculture (Figure 6E). Furthermore, the level of expression was similar with human primary blood capillary endothelial cells. Therefore, the possibility that endothelial cells that we observed in the presence of MDM are of lymphatic nature is unlikely.

In chick kidney development, stromal compartment and partially tubular capillary originate from paraxial mesoderm^42^. Furthermore, using three lineage tracing methods, Peng et al. showed recently that even nephron progenitors can originate from somites in Zebrafish^43^. Somites-derived cells were detected in glomeruli of adult zebrafish. In human developing kidney transcriptional profile of nephron progenitors and interstitial progenitors have overlapped significantly and FoxD1 expression was detected in Six2-positive nephron progenitor cells^44^. Hence, it is possible that improved development of organoids in the presence of MDM is due to strongly increased development of paraxial mesoderm and somitogenesis. Precise mechanisms of paraxial mesoderm induction by MDM are still being investigated by us.

In addition, we have detected strong induction of FoxD1 gene expression on day 8. Furthermore, we detected a population of FoxD1/CD31 double-positive cells on day 8 and day 14. Stromal compartment and FoxD1 in particular, are important for kidney development. Thus, FoxD1-knockout mice demonstrate kidney hypodysplasia, fused kidney and pelvic location, fused glomeruli and delayed nephrogenesis^28^. Those malformations result from disrupted interstitium and changed composition and patterning of extracellular matrix in the kidney. By means of human proteome profiler array we have detected expression of several cytokines including ENA-78 (CXCL5) by MDM (Figure 3). ENA-78 has been implicated in the induction of FoxD1 expression in cancer cells^30^. In differentiating kidney organoids, exogenously added ENA-78 strongly induced FoxD1 expression in the absence of MDM (Figure 7). Thus, MDM-released ENA-78 induces expression of FoxD1, that results in endothelial differentiation and promotion of organoid development.

Our organoid model has several important advantages over those reported earlier. First, we demonstrate development of vasculature in kidney organoids that was not possible to achieve without genetic modifications of iPSC. Second, we discovered novel mechanisms of endothelial cell differentiation and vascularization in renal organoids. Our model provides an opportunity to investigate developmental mechanisms that were not previously observed in vitro. Third, the increased vascularization provides a basis for further improvements of organoid development in vitro by tissue engineering approaches. Most likely, our observed effects are not limited to kidney organoid development since absence of vascularization is a common problem of the organoid field.

## Material and methods

The methods and protocols employed adhered to pertinent guidelines and regulations and received approval from the Ethics Committee of the Hannover Medical School. Biological safety levels S1 and S2 were implemented according to standard practices. All information regarding the chemicals, kits, equipment, composition of chemicals and consumables, as well as the formulation and preparation of buffers and media utilized, is provided in detail and elaborated upon in the concluding section of the chapter.

### Cells and cell culture

During the study, the following cell types were used: Epi-iPSC (Gibco by ThermoFisher, A18945), ED-iPSC, which was bestowed upon us as a gift by Dr. Aloise Mabondzo (CEA, Institute Joliot, Paris, France)^45^ cl11/Selene CD34+ cord-blood derived iPSC was a kind gift of Dr. Nico Lachmann (Hannover Medical School, Hannover, Germany) (https://hpscreg.eu/cell-line/MHHi015-B). iPSCs were maintained in StemFlex medium (Gibco by ThermoFisher, A3349401) at 37°C with 5% CO2, as recommended. StemFlex medium was replaced every 2 days, and upon reaching 90% confluence, cells were split at a 1:3 ratio using StemPro Accutase (Gibco by ThermoFisher, A1110501). CD31 promoter - Gluc iPSC reporter cell line, which expresses secreted Gaussia luciferase under the control of the CD31 promoter (CD31-Gluc iPSC), and a cell line with the constitutive promoter of human origin EF1alpha as a negative control for the CD31-Gluc iPSC cell line (NC-iPSC) were established in our laboratory from Epi-iPSC. Human leukemia monocytic THP-1 cell line (THP-1) (Leibniz Institute DSMZ German Collection of Microorganisms and Cell Cultures GmbH, ACC 16) was maintained in RPMI 1640 medium (PAN-Biotech, P04-16500) supplemented with 20% Fetal Bovine Serum (FBS) (PAN-Biotech, P30-3306) and 1% Penicillin-Streptomycin (PAN-Biotech, P06-07100). Human Embryonic Kidney cell line (HEK 293, ATCC CRL-1573) was maintained in DMEM medium (Gibco by ThermoFisher, 11965092) supplemented with 10% Fetal Bovine Serum (FBS) (PAN-Biotech, P30-3306) and 1% Penicillin-Streptomycin (PAN-Biotech, P06-07100). Human iPSC-macrophages, kindly provided to us as a gift by Prof. Dr. Nico Lachmann (Hannover Medical School, Hannover, Germany) (DOI: 10.1016/j.stemcr.2015.01.005, DOI: 10.1038/s41596-021-00654-7). Peripheral blood classical CD14+/CD16-monocytes were isolated from buffy coats provided by the German Red Cross (DRK-Blutspendedienst NSTOB, Springe, Germany). Informed consent was obtained from all donors. Monocytes were isolated using the Classical Monocyte Isolation Kit (Miltenyi Biotec, 130-117-337) according to the manufacturer’s instructions.

To develop a CD31 promoter - Gluc iPSC reporter cell line, we used a lentiviral Human PECAM1 promoter reporter clone from GeneCopoeia, Inc. (LPP-HPRM30418-LvPG04-A00). Briefly, HEK 293T cells were transfected with Packaging Plasmid (psPAX2), Envelope Plasmid (pMD2.G) and transfer plasmid using Perfectin transfection reagent from Genlantis (T303007). Subsequently, Lentiviral particles were concentrated by ultracentrifugation and used to infect Epi-iPSCs. Puromycin selection was then performed to obtain a CD31-Gluc stable cell line. For normalization of Gaussia luciferase expression, we generated a control iPSC - cell line with an irrelevant constitutive promoter of human origin EF1alpha as a negative control. To measure luciferase activity, Pierce Gaussia Luciferase Glow Assay Kit (ThermoFisher, 16161) was used as recommended by the supplier. Luminescence was then assessed using a microplate reader TECAN Infinite 200 PRO equipped with a 450 nm filter, with an integration time of 1000 ms, gain of 100, and Relative Light Units (RLU) of 100 or 150.

To isolate CD31+ cells, a digestion mixture was first developed to generate a single-cell suspension from three-dimensional organoid structures. The mixture contained 17% of 13 U/mL Liberase (Roche by Merck, 5401020001), 0.2% DNase (QIAGEN, 79254), and 14% Hyaluronidase (1mg/mL in H2O, Sigma-Aldrich by Merck, 37326-33-3) in DPBS. The organoids were washed with warm DPBS, and 700µL of the digestion mix was added per well in a 6-well plate, then checked after incubating for 20 minutes at 37°C. The single-cell suspension was filtered, and Dynabeads CD31 Endothelial Cell (Invitrogen by ThermoFisher, 01261441) were used to isolate CD31+ cells following the manufacturer’s instructions.

iPSC cultivation for macrophages differentiation was performed using a feeder-free protocol^46^. Briefly, iPSC cells were cultured on Geltrex-coated tissue culture plates or T25 flasks using E8 medium (containing DMEM/F-12 (Gibco, Life Technologies), 64 mg/L ascorbic acid 2-phosphate, 14 µg/L sodium selenite, 543 mg/L, NaHCO3 and 20 mg/L insulin, and 10.7 mg/L human recombinant transferrin (all from Sigma-Aldrich) supplemented with 100 ng/mL human basic fibroblast growth factor (hbFGF) and 2 ng/mL human transforming growth factor beta (TGFß) (both from PeproTech) under standard humidified conditions at 37°C and 5% CO2. The cells were split two times a week using Accutase (STEMCELL Technologies) and under the addition of 10 µM ROCK inhibitor (RI; Tocris, Bristol, UK). A complete medium change using ROCK inhibitor-free medium was performed 48 hours after splitting. In order to start mesoderm priming, 5×10^5^ iPSC were seeded in 3 mL of mesoderm priming I medium (E8 medium supplemented with 10 µM RI, 50 ng/mL human vascular endothelial growth factor (hVEGF) and human bone morphogenetic protein 4 (hBMP4) and 20 ng/mL hSCF (all from PeproTech) using CELLSTAR 6-well plates placed on an orbital shaker at 70 rpm, leading to the formation of embryoid bodies (EBs). On day 2, the medium was changed to E6 medium (containing only hVEGF, hBMP4 and hSCF). On day 4 after mesoderm priming I, supernatant was discarded and 3 mL of mesoderm priming II medium (E6 medium supplemented with 50 ng/mL hVEGF and hBMP4, 20 ng/mL hSCF and 25 ng/mL human interleukin 3 (hIL-3)) was added, while increasing the shaking speed to 85 rpm. Mesoderm priming medium II was refreshed at day 7 before macrophage differentiation (hematopoietic differentiation) was started at day 10. For this, the medium was removed and the EBs were transferred to a 6-well tissue culture plate using 2 mL of differentiation medium (X-VIVO containing 1% (v/v) P/S and L-Glu, 0.1% (v/v) β-mercaptoethanol supplemented with 25 ng/mL hIL-3 and 50 ng/mL human macrophage colony stimulating factor (hM-CSF). Macrophage production started from day 4 onwards and can be continuously harvested 1–2×per week. The harvested macrophages were filtered (70 µm pore size) and seeded in Roswell Park Memorial Institute (RPMI 1640) medium containing 1% (v/v) P/S, 10% (v/v) fetal bovine serum (FBS) and 50 ng/mL hM-CSF using tissue culture plates.

### Human iPSC-derived kidney organoids differentiation

KORGs differentiation was conducted following the protocol outlined by Morizane et al.^47^, with some modifications. Completely confluent iPSC were split at a 1:2 ratio the day before differentiation, as described above. For co-culturing with monocytes, THP-1 and iPSC-derived macrophages, we used 24-well or 6-well plates with transparent Thincert cell culture inserts, featuring a pore diameter of 0.4 µm (Greiner Bio-One, 657641 or 662641). Monocytic cells were seeded at a density of 1 × 10^5^ or 2 × 10^5^ cells, depending on the size of the plate. Medium was replenished in the same manner as during KORGs differentiation. Mono-cultured myeloid cells were also placed in the inserts and treated in the same way without iPSC.

Advanced RPMI 1640 Medium (Gibco by ThermoFisher, 12633012) supplemented with 200 µM L-Glutamine (PAN-Biotech, P04-80100) and 0.5% KnockOut Serum Replacement (KOSR) was used as the BM for kidney organoids. To induce late primitive streak induction on the first day of differentiation, 10 µM Laduviglusib (CHIR-99021) HCl (Selleckchem, S2924) and 5 ng/mL Recombinant Human Noggin (PeproTech by ThermoFisher, 120-10C) were added. The medium was refreshed after two days. From the fourth to the seventh day, the BM was enriched with 10 ng/mL Human/Mouse/Rat Activin A Recombinant Protein (E. coli derived, PeproTech by ThermoFisher, 120-14E), inducing posterior intermediate mesoderm induction. Subsequently, for the next two days, cells were differentiated in BM with 10 ng/mL Recombinant Human FGF-9 Protein (R&D Systems, Inc., a Bio-Techne Brand, 273-F9-025). From the ninth to the eleventh day, organoid progenitors were supplemented with BM and an additional 3 µM CHIR-99021, maintaining the same concentration of FGF-9. During the following three days, BM was supplemented with 10 ng/mL FGF-9 only, inducing nephron progenitor cell activation. In the last part of the protocol, which lasted from the fourteenth to the twenty-eighth day, only BM was used for the cells, with refreshing every two days.

### KORGs fixation and staining

To proceed with fluorescence microscopy, we followed the protocol outlined below. Initially, 4% paraformaldehyde in DPBS was applied for 60 minutes at room temperature to fix the organoids. Subsequently, the KORGs underwent three washes using 0.1% Triton X-100 in DPBS. To block nonspecific binding, a solution consisting of 5% goat serum, 5% donkey serum, 1% Triton X-100, and 5% BSA in DPBS was added for 60 minutes twice. Following this, the organoids were left to incubate at 4°C overnight with primary antibodies. For the addition of secondary antibodies, the KORGs underwent three washes in 1%Triton X-100 in DPBS for 60 minutes each time. The secondary antibodies were then added for overnight incubation at 4°C or at room temperature for 3 hours. After the incubation with secondary antibodies, the organoids were exposed to a series of Formamide/PBS solutions (vol/vol): 20% for half an hour, 40% for half an hour, 80% for one hour, 95% for half an hour. Finally, the organoids were cleared using a 95% Formamide (Sigma, SLCH8231) in PBS solution overnight and were mounted with Aqua-Poly/Mount (Polysciences, 18606-20) the next day. Fluorescence microscopy was conducted at the Research Core Unit for Laser Microscopy at Hannover Medical School, utilizing an oil-immersed x20 objective for image capture. The obtained images were subsequently analyzed using ImageJ analysis software.

### Imaging and quantitative analysis of vasculature

Confocal microscopy was performed using Leica TCS-SP2 AOBS confocal microscope (Leica Microsystems) at the Research Core Unit for Laser Microscopy at Hannover Medical School. Processing of the 3D confocal stacks was performed using ImageJ analysis software (ImageJ 1.48v; https://imagej.nih.gov/ij/). Vascular network was quantitatively analyzed by AngioTool64 software^19^ AF488-CD31 channel was selected and images exported in Tag Image File Format (TIFF). The μm size of the original scanned area was recorded to standardize all samples. The low threshold was adjusted for each individual image between 10 and 29, with co-culture images having a higher low threshold. Small Particle threshold was also individually adjusted between 0 and 800, and fill holes between 0 and 600. Scaling factor remained at 1 for all images. Some parameters were standardised between samples using the μm size of their respective original scanned area. Main Analysed parameters were: Average vessel length, total number of endpoints/μm (open-ended segments), total number of junctions/μm (vessel branching), total vessel length/μm, vessel area/μm, vessel percentage area and mean endothelial lacunarity (measure of fractals and spatial arrangements whereas higher number of gaps leads to higher mean E. lacunarity which means less developed and more chaotic endothelial networks).

### Flow cytometry

Peripheral blood mononuclear myeloid cells (PBMMC) in inserts were suspended in FACS buffer consisting of DPBS (Sigma-Aldrich), 2% fetal calf serum (FCS) (Biochrom), 2 mM Na_2_EDTA (Roth), and 0.02% NaN_3_ (AppliChem). Human TrueStain FcX^TM^ antibody (hFc) was used to block nonspecific binding. Zombie R718^TM^ Fixable Viability Kit was used to stain dead cells. PBMMC were stained with the anti-human extracellular antibodies specific to MerTK (MerTK), CD45 (PTPRC), CD163 (M130), CD16 (FCγR3A), CD14 (CD14), HLA-DR (HLA-DR), CD11c (ITGAX), CD86 (B70) and anti-human intracellular antibodies specific to CD68 (LAMP4) and CD206 (MMR). All antibodies are from BioLegend. Antibody dilutions are listed in the Supplementary Table 4. Exposure time for intra and extra-cellular antibodies was 25 min. hFc staining was 15 min and Zombie was 30 min. All cell staining, except Zombie (room temperature and dark), was performed on ice and in the dark. Extracellular staining was performed using FACS buffer, intracellular staining was performed using BD Cytofix/Cytoperm™ Fixation/Permeabilization Kit (BD Biosciences) and BD Perm/Wash™ (BD Biosciences), and Alive/dead cell staining was performed with DPBS. Single stain compensations were performed for every sample. Acquisition was performed on an LSR-II flow cytometer (BD Biosciences). FACS data was analyzed using FlowJo software (FlowJo LLC). Doublets were identified and excluded based on forward (FSC) and side (SSC) scatter parameters. Dead cells (Zombie 10^4^) were excluded. Analysed myeloid cells were identified using CD45+ gate and CD14+ gate.

### RNA purification and RT-PCR

Qiagen RNeasy Mini Kit (Qiagen, 74106) was used for RNA purification for RT-PCR assay. RNeasy Plus Mini Kit (Qiagen, 74136) were utilized to extract RNA for bulk RNA-sequencing. Each RNA sample was derived from approximately 100 organoids. Concentration and quality of RNA were assessed using the NanoPhotometer® N60 (Implen). TaqMan verified gene expression assays used in the study are listed in the supplementary table X. TaqMan RT-PCR was performed using Applied Biosystems TaqMan RNA-to-CT1-Step Kit (ThermoFisher 4392938) as recommended.

### RNA-sequencing analysis

After determining RNA integrity and quality, RNA was prepared and sequenced at Eurofins Genomics (Constance, Germany) using an INVIEW Transcriptome (NGS Built For You (eurofinsgenomics.eu)) product. This included purification of mRNA, fragmentation, strand-specific cDNA synthesis, end-repair, ligation of sequencing adapters, amplification and purification. The prepared libraries were then quality-checked, pooled and sequenced on an Illumina platform (Illumina NovaSeq X+, PE150 mode). Bioinformatic analyses were performed using the open-source Galaxy platform (version 25.0.3.dev0). The tool Falco was used for initial quality control checks. Futher Cutadapt was applied with a quality cutoff of 20 and a minimum length of 20 as the main filtering parameters. This step was used to identify and remove adapter sequences, primers, poly-A tails, and other types of unwanted sequences from the reads. Reads were mapped to the human genome (hg38 | GRCh38) using STAR, and gene-level counts were obtained with featureCounts. For differential gene expression (DGE) analysis, DESeq2 was used. Since we had only one experimental factor with three biological replicates, the basic DESeq2 setup was sufficient. Heatmaps were generated using the heatmap2 tool in Galaxy, and volcano plots were created using the Volcano Plot tool. To highlight biological processes, we applied goseq and pathview for Gene Ontology (GO) analysis of RNA-seq data and for generating KEGG pathway visualizations of differentially expressed genes. Genes were considered differentially expressed if they had an adjusted *p*-value < 0.05 and a log2 fold change > 1 or < –1.

### Statistics and Graphs

All data were obtained with at least 3 biological replications. Data are presented as mean ± standard deviation (SD). Multiple comparisons were analyzed by ANOVA with Tukey *post hoc* test. P-values < 0.05 were considered statistically significant. GraphPad Prism 8.3.0 (GraphPad Software) and GraphPad prism 9 were used for data analysis. FACS data analysis was performed using Two-way Anova. Kruskal-Wallis test was applied for AngioTool image analysis.

## Acknowledgements

This work was supported by the Deutsche Forschungsgemeinschaft (DFG, German Research Foundation) under Germany’s Excellence Strategy (EXC 2155; RESIST; project number 390874280 and DFG support LA 3680/9-1 and 10-1) (N.L.); the REBIRTH Research Center for Translational Regenerative Medicine ‘Förderung aus Mitteln des Niedersächsischen Vorab’ (Grant ZN3340) (N.L.); the European Research Council (ERC) under the European Union (EU)’s Horizon 2020 research and innovation program (Grant agreement 852178); and the EU (Grant agreement 101100859 and 101158172) (N.L.). The views and opinions expressed are those of the authors only and do not necessarily reflect those of the EU or the ERC. Neither the EU nor the granting authority can be held responsible for them. The project was additionally supported by zukunft.niedersachsen (Federal State of Lower Saxony), R2N.Micro-Replace-Systems. Additional funding was provided by the German Center of Lung Research (DZL) and the Federal Ministry of Research, Technology and Space (BMFTR, SMARTibone project). This work was supported by the Fraunhofer Internal Programs under Grant No. Attract 40-01696. In addition, was supported by the Deutsche Forschungsgemeinschaft (DFG, German Research Foundation) grant KA 5549/2-1.

