## Supplementary figures for "Macrophages induce stromal differentiation and endothelialization in iPSC-derived kidney organoids"

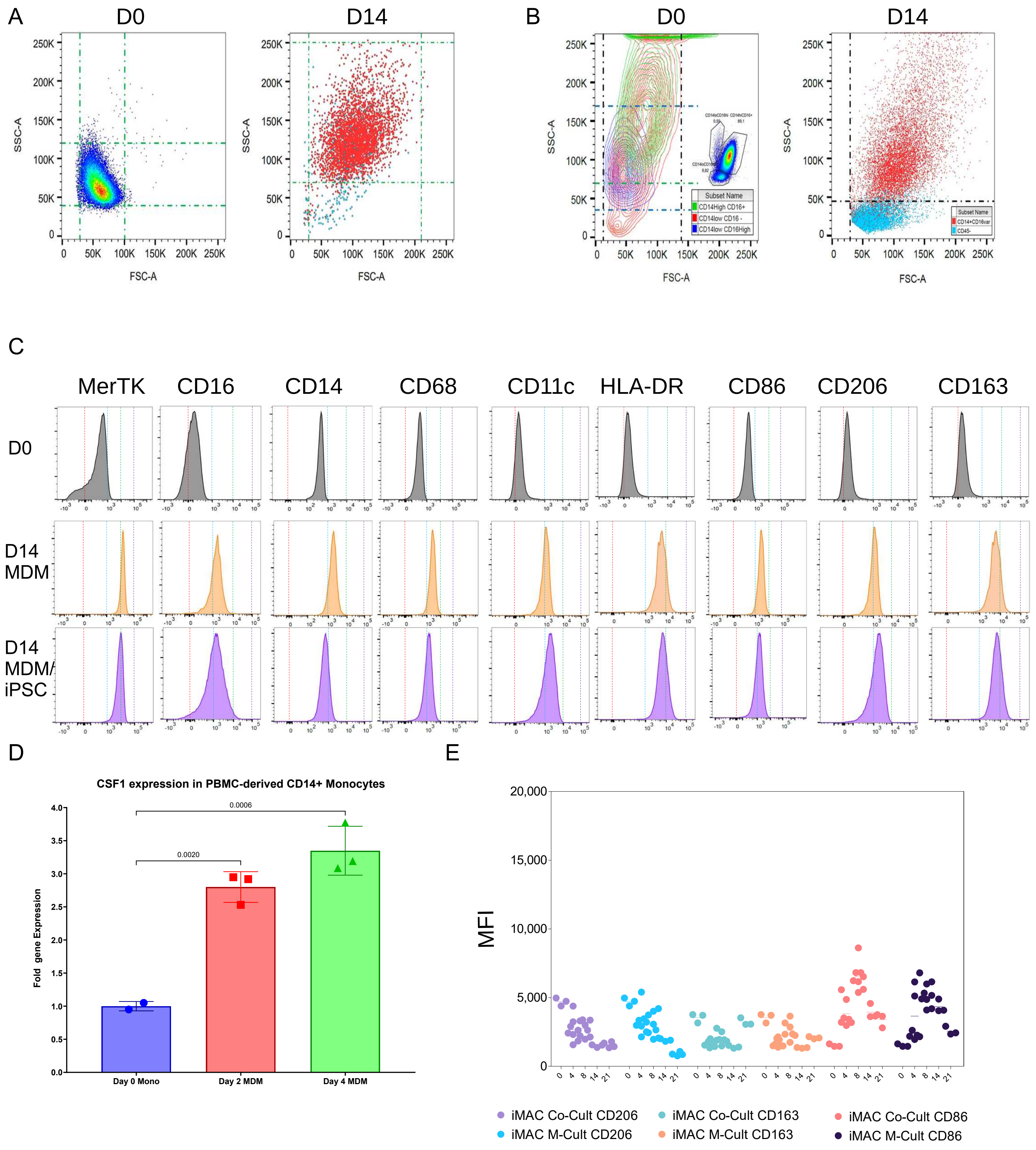


**Supplementary figure 1.** A-B. Morphology of MDM (A) and iPSC-derived macrophages (B) on day 14 co-culture with iPSC was assessed based on FSC and SSC parameters using FACS. C. Expression of macrophage markers in MDM on day 0 and day 14 of exposure to kidney organoid differentiation factors, with or without iPSC. D. M-CSDF expression by monocytes exposed to to kidney organoid differentiation factors on day 2 and day 4. E. IPSC-derived macrophages showed less changes in markers expression than MDM.

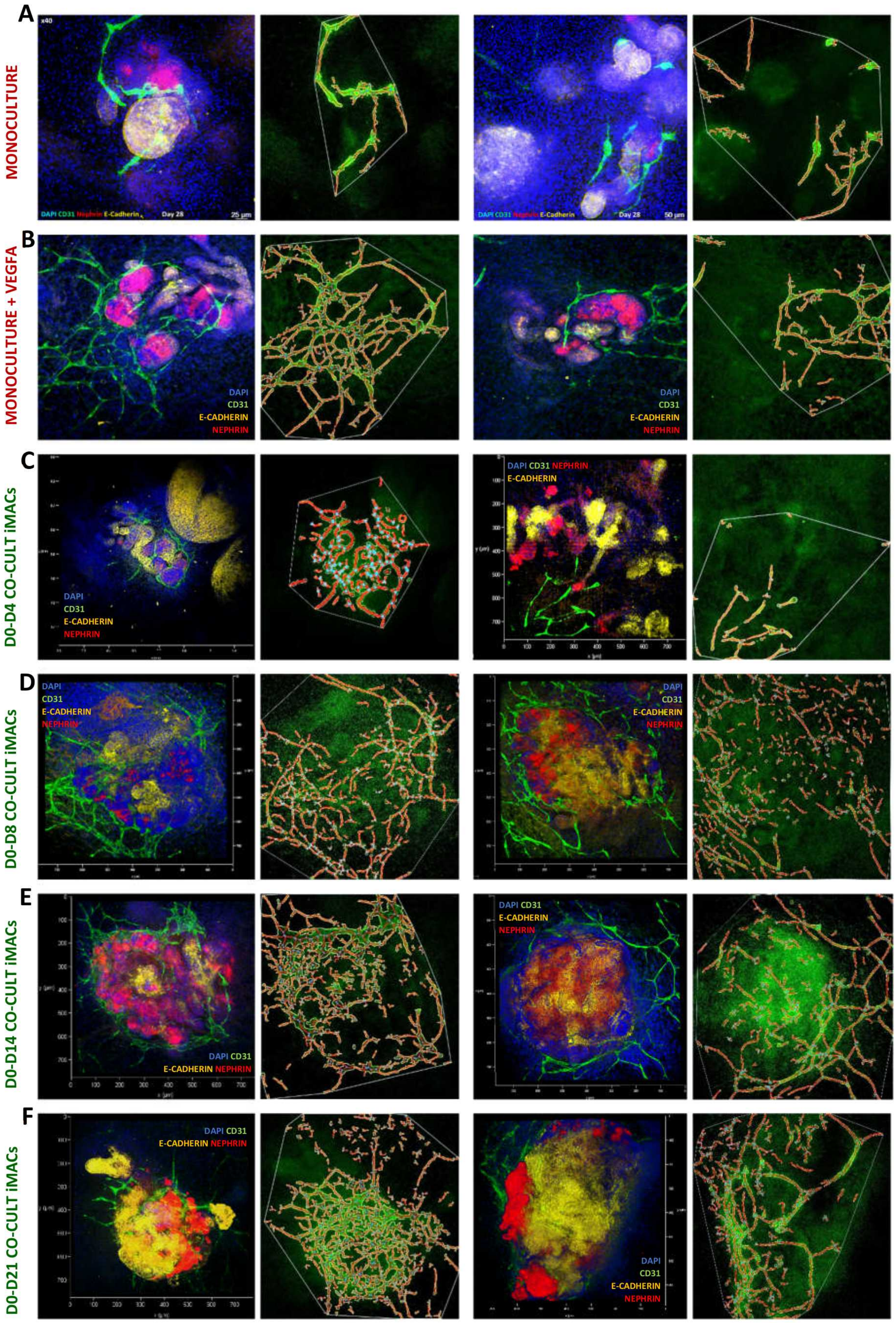


**Supplementary Figure 2.** Organoids from cl11/Selene-iPSC in mono-culture and co-culture with iiPSC-derived macophages were stained on day 28 for AngioTool analysis. Examples of trimmed images of CD31+ vasculature parameters detected by AngioTool. Parameters: vessel area, length, percentage, junctions, density, end points and lacunarity.

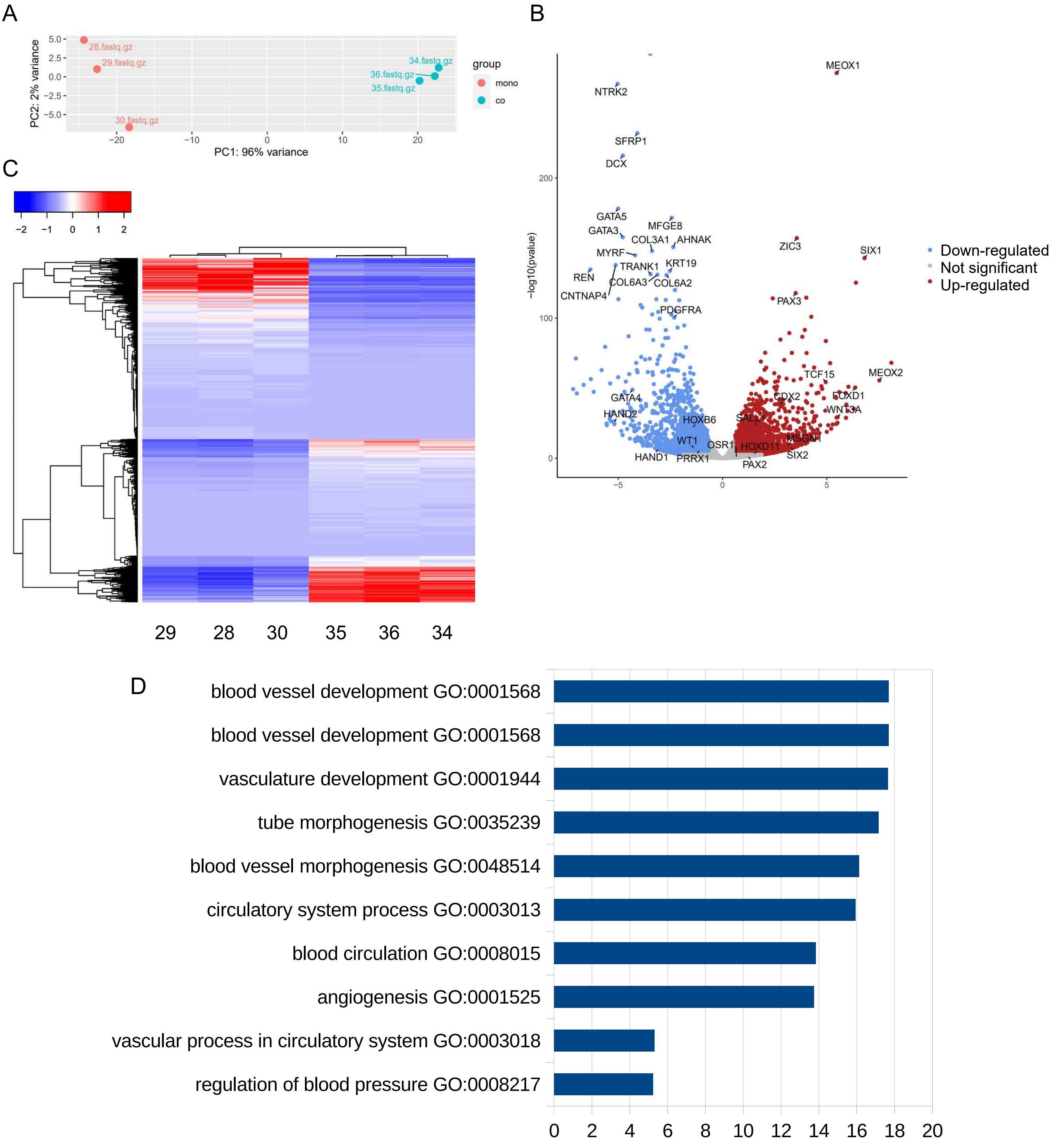


**Supplementary figure 3.** Bulk RNA-sequencing analysis of Epi-iPSC co-cultured with MDM versus mono-cultured Epi-iPSC. A. PCA plot. B. Volcano plot. C. GO identified by enrichment analysis of downregulated DEG.

A

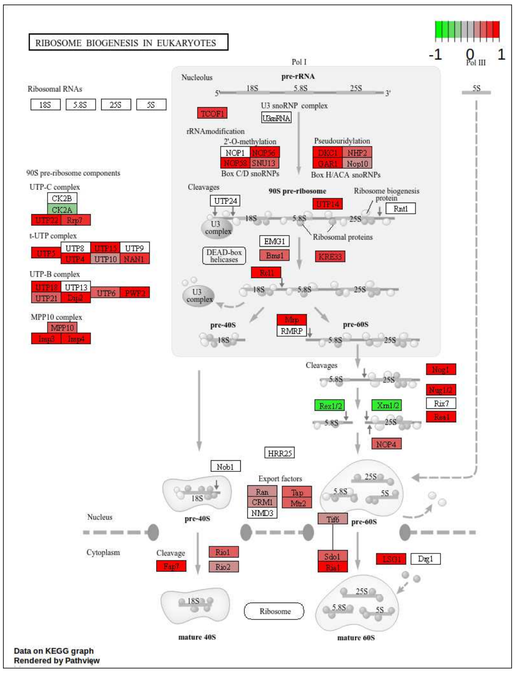


B

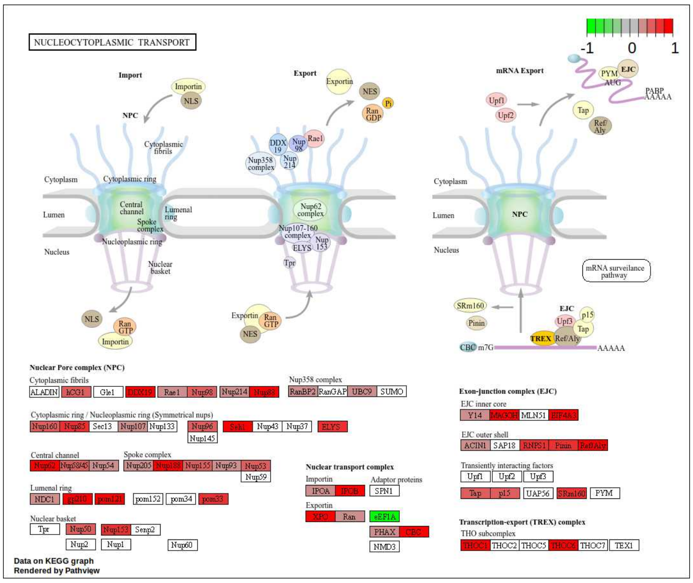


C

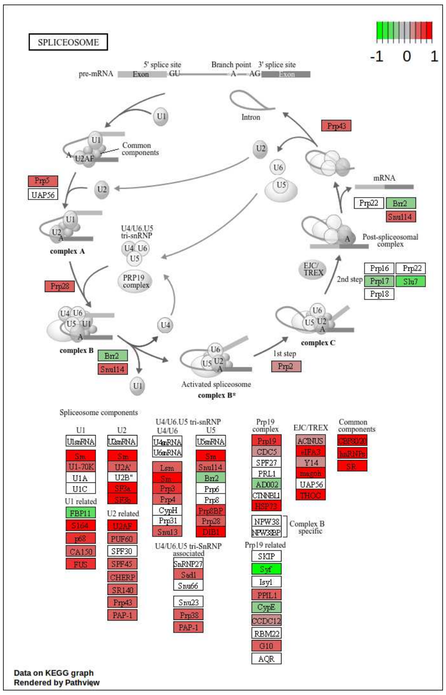


**Supplementary figure 4.** Visualization of some signaling pathways identified by KEGG enrichment analysis of Epi-iPSC co-cultured with MDM versus mono-cultured Epi-iPSC with gene expression values mapped to gradient color scale.
